# Contact Stiffness Provides a Unified Frame of Reference for Understanding the Effects of Extracellular Matrix Mechanics on Cell Behaviors

**DOI:** 10.1101/2022.09.07.506641

**Authors:** Peng Zhao, Zhaoyi Zhang, Yang Zheng, Yina Gao, Jialing Cao, Mingwei Jiang, Yuxuan Jiang, Li Gao, Jing Du, Yanping Cao

## Abstract

In interactions between cells and extracellular matrices (ECMs), contact mechanics theory indicates that local ECM deformation depends on both local and non-local forces imposed by cells. In the present study, we investigated the use of a comprehensive variable, contact stiffness (CS), to interpret cell-ECM interactions. CS defines the relationship between the local ECM deformation and the total force from a cell, integrating the effects of individual variables including ECM stiffness, ECM thickness, and cell adhesion area. Through assessments of ECM mechanosensing by human mesenchymal stem cells (hMSCs) under varied CS conditions, we showed that CS scaled well with both yes-associated protein (YAP) activity and the extent of stem cell differentiation. To reveal the cross-scale mechanism underlying mechanosensing, we propose a CS-based motor clutch model, which suggests that various mechanical stimuli affect cells by altering the CS, thus altering the reaction force from the ECM. Using the proposed model, we revealed the contributions of cell architecture evolution to stem cell differentiation and predicted the influence of a non-adjacent ECM layer on cellular mechanosensing. These results demonstrate that the use of CS provides a quantitative predictive framework that allows researchers to address longstanding questions about the effects of ECM mechanics on cell behaviors.

## INTRODUCTION

When cells interact with the extracellular matrix (ECM), they are continuously exposed to physical forces imposed by the surrounding microenvironment, and respond by modulating their behaviors and generating their own forces (*1–3*). Emerging evidence shows that mechanical variables in the cell-ECM system, including ECM stiffness (*4–6*), ECM thickness (*7, 8*), and cell adhesion area (*9, 10*), have profound effects on cell behaviors (*e.g.*, motility (*11–13*), proliferation (*14*), apoptosis (*15*), and differentiation (*6, 16, 17*)) that are required for processes such as embryonic development (*12*), tissue regeneration (*18*), and tumor metastasis (*19, 20*). For example, mesoderm stiffening is essential to triggering neural crest cell migration in *Xenopus laevis* (*12*), and matrix stiffness promotes tumor metastasis by driving the epithelial-mesenchymal transition of breast tumor cells (*5*).

Cells sense and response to mechanical stimuli arising from ECMs through cycles consisting of mechanosensing followed by mechanotransduction, then mechanoresponse (*2*). The mechanisms underlying ECM mechanosensing have been explained with motor clutch models, which suggest that cells perceive alterations in ECM mechanics through an intricate balance between pulling forces, which are actuated by actin fibers on focal adhesion (FA) plaques, and mechanical reactions of the ECM (*21*). Importantly, previous models of ECM mechanosensing essentially assume that the local reaction force of the ECM on the cell depends on the local ECM deformation state and ECM stiffness. However, contact mechanics theory (*22*) has established that local ECM deformation depends on both local forces (exerted by local FAs) and non-local forces (imposed by the cell through non-local FAs); *i.e.*, variations in local ECM deformation are induced by changes in the total force, rather than the local force, imposed by the cell on the ECM. Therefore, the local reaction force of the ECM on the cell cannot be quantified solely from the local ECM deformation and stiffness. Such a deformation-force relationship would depend simultaneously on several ECM-associated variables (including ECM stiffness and thickness) and variables related to cell-ECM contact geometry (including adhesion area and shape). Previous studies focusing on these variables individually have advanced our understanding of how ECM mechanosensing and associated cellular processes regulate diverse physiological and pathological processes (*4–10*). However, contact mechanics theory indicates that the effects of these variables are not mutually independent (*22*), but rather are coupled in the mechanosensing process (*23*). The underlying mechanisms by which these different variables synergistically regulate cell behaviors remain unknown.

To address this issue, we here investigated contact stiffness (CS), which defines the relationship between local ECM deformation and the force from a cell, in the context of the mechanical effects of the ECM on cell behaviors. Based on contact mechanics theory (*22*), CS integrates the effects of individual parameters (including the stiffness and thickness of the ECM and the area and shape of cell adhesion) into one variable that impacts cell-ECM interactions. We assessed ECM mechanosensing in human mesenchymal stem cells (hMSCs) under conditions with varying CS values. The results demonstrated that CS outperformed other mechanical variables (*e.g.*, ECM stiffness, cell adhesion area, and ECM thickness) in characterizing mechanoregulation of YAP activation and stem cell differentiation. Using the CS, we here propose a cross-scale model based on the classical motor clutch model, which reveals the contributions of cell architecture to stem cell differentiation and predicts cellular mechanosensing of a non-adjacent ECM layer. Our findings demonstrate that CS provides a unified frame of reference for understanding the mechanosensing processes by which cells respond to various mechanical stimuli from ECMs.

## RESULTS

### CS defines the relationship between local ECM deformation and the force imposed by a cell

Contact mechanics theory (*22, 24, 25*) indicates that during cell-ECM interactions, the variation in local ECM deformation (*D*) is related to changes in the total force (*F*) from the cell by the CS (*S*), *i.e.*, d *F* = *S* d *D*. CS is a variable that depends on the mechanical variables of the ECM, including ECM stiffness (as measured by the elastic modulus and Poisson’s ratio for an elastic ECM), ECM thickness, and the geometric variables defining cell-ECM contact geometry, including the cell adhesion area and shape. In cases where the ECM is far thicker than a cell (> 10-fold), CS can be expressed 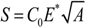 (*26*), where *C*_0_ is a constant; *E\** is the plane-strain modulus (which defines ECM stiffness); and *A* is the cell adhesion area (*26*). When the ECM thickness must be considered (*i.e.*, ECM thickness is on the same order as the dimensions of the cell), CS can be expressed as 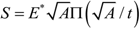 where Π is a dimensionless function and *t* is the ECM thickness (Fig. 1A).

**Fig. 1.**
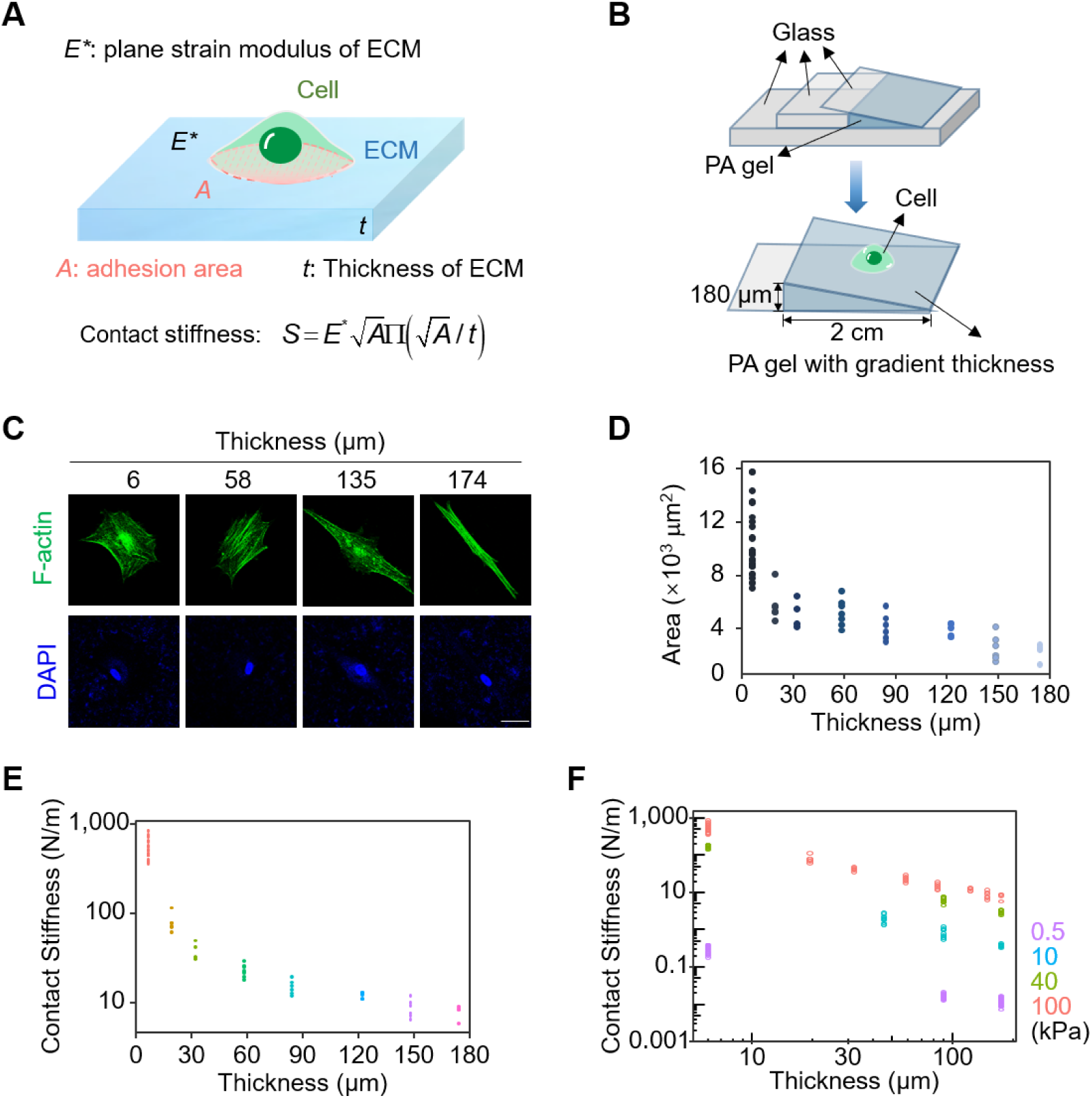
Contact stiffness defines the relationship between local ECM deformation and the force imposed by the cell. (**A**) Schematic diagram of the definition and calculation of contact stiffness during cell-ECM interactions. (**B**) Schematic diagram of the fabrication of ECM composed with a gradient thickness gel. Since the gradient of gel thickness is rather small, the variation in gel thickness across a cell dimension is negligible. (**C**) Representative images of F-actin and nucleus in hMSCs grown on a 100 kPa gradient thickness gel at distinct thickness regions stained by phalloidin (green) and DAPI (blue), respectively. Scale bar: 30 μm. (**D**) The statistical analysis of cell adhesion area in hMSCs grown on a 100 kPa gradient thickness gel at distinct thickness regions as shown in (C). (**E**) The statistical analysis of contact stiffness in hMSCs grown on a 100 kPa gradient thickness gel at distinct thickness regions as shown in (C). (**F**) The statistical analysis of contact stiffness of hMSCs grown on gradient thickness gels with different stiffness (0.5, 10, 40, and 100 kPa) at distinct thickness regions.

Because CS defines the relationship between local ECM deformation and the cell-ECM interaction force and integrates individual variables such as ECM stiffness, ECM thickness, and cell adhesion area and shape into one variable, we examined the use of CS to investigate ECM mechanosensing. To avoid the influence of chemical factors in preparing ECMs with a wide range of CS values, we cultured mechanosensitive hMSCs on a layered ECM composed of polyacrylamide (PA) gel in a gradient of thicknesses on a glass slip (here referred to as a gradient thickness gel) (Fig. 1B). Although the gel stiffness was consistent across the gradient, the cell adhesion area increased as the gel thickness decreased. This was demonstrated with immunofluorescent staining of integrin and F-actin; the adhesion area increased from 1,000 to 16,000 μm^2^ as the gel thickness decreased from 174–6 μm on a 100-kPa gel (Fig. 1C, D; Supplementary Fig. S1). In this system, CS varied as a function of gel thickness. Specifically, CS increased from 8–547 N/m as the gel thickness decreased from 174–6 μm on a 100-kPa gel (Fig. 1E).

To achieve a wider range of variation in CS, we combined datasets from gradient thickness gels with different stiffnesses (namely 0.5, 10, 40, and 100 kPa). In the combined dataset, the CS ranged over five orders of magnitude from 0.012–547 N/m, which was much larger than the range of variation in ECM stiffness. This broad range of CS variations and its physical meaning indicated that CS is more informative in this context than individual variables. Furthermore, previous studies of cell-ECM interactions in which individual variables (such as ECM stiffness, ECM thickness, or the cell adhesion area) were tuned also modulated the CS experienced by a cell. CS therefore represents a promising mechanical variable for interpreting the integrated effects of these individual variables on ECM mechanosensing.

### CS outperforms other individual variables of cell-ECM systems in interpreting mechanical regulation of YAP

Emerging evidence has revealed that YAP activity is regulated by various mechanical variables such as ECM stiffness, ECM thickness, and cell adhesion area (*9, 23, 27, 28*). To reveal the coupled effects of these individual variables, we explored the use of CS for interpreting mechanical regulation of YAP activation. We calculated YAP nuclear translocalization in terms of the nuclear/cytoplasmic distribution (N/C ratio) in hMSCs cultured on gradient thickness gels. Immunofluorescent staining showed that the YAP N/C ratio continuously increased along with increases in CS as the gel stiffness remained constant (Fig. 2A, B). Remarkably, a comparison of cells grown on gradient thickness gels with different stiffnesses showed that the YAP N/C ratio was similar when CS values were comparable despite vast differences in either gel stiffness or thickness. For example, YAP N/C ratios were comparable between a 10-kPa ECM that was 135 μm thick (CS = 3.7 N/m) and a 40-kPa ECM that was 40 μm thick (CS = 3.1 N/m) (Fig. 2C, D). These results indicated that cells sensed CS rather than ECM stiffness when activating YAP. Thus, exploring the mechanosensing capacities associated with YAP activation in terms of CS instead of ECM stiffness will likely yield more informative inferences.

**Fig. 2.**
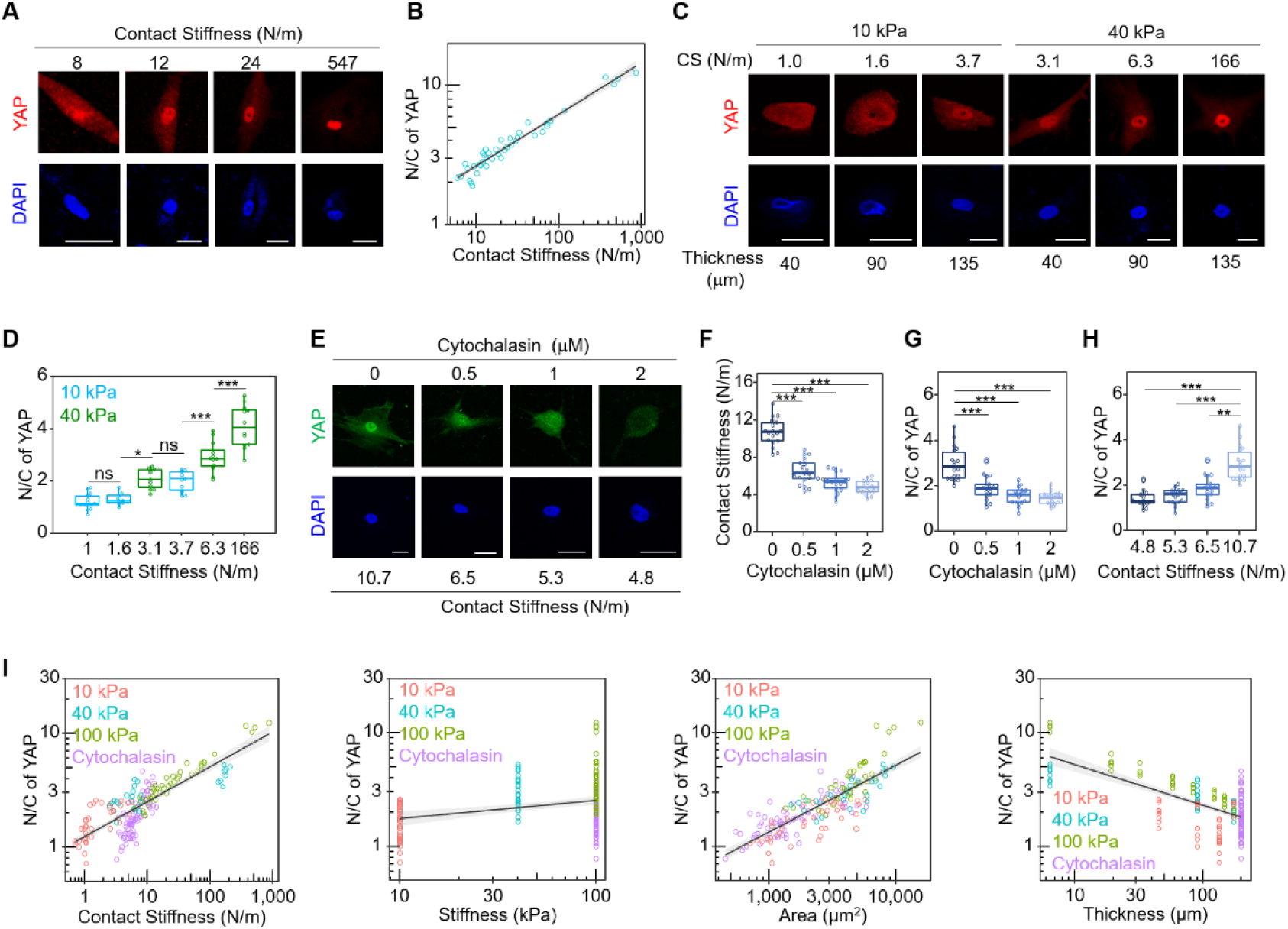
Contact stiffness outperforms other individual parameters of cell-ECM systems in interpreting the mechanical regulation of YAP. (**A**) Representative immunofluorescence images of YAP and nucleus in hMSCs grown on a 100 kPa gradient thickness gel at different contact stiffness stained by YAP antibody (red) and DAPI (blue), respectively. Scale bar: 30 μm. (**B**) The statistical analysis of YAP N/C ratio in hMSCs grown on a 100 kPa gradient thickness gel at different contact stiffness as shown in (a). YAP N/C ratio scales as power functions of Contact Stiffness, with exponent 0.357 ± 0.009 (*r^2^* = 0.967, *P* < 0.001). Grey area are 95% confidence intervals for the fitted functions. (**C**) Representative immunofluorescence images of YAP and nucleus in hMSCs grown on a 10 or 40 kPa gradient thickness gel at different contact stiffness stained by YAP antibody (red) and DAPI (blue), respectively, Scale bar: 30 μm. (**D**) The statistical analysis of YAP N/C ratio in hMSCs grown on a 10 or 40 kPa gradient thickness gel at different contact stiffness as shown in (C). All error bars are S.E.M. (n = 14, 9, 10, 10, 12, and 12 cells for CS = 1, 1.6, 3.1, 3.7, 6.3, and 166 N/m, respectively. \**P* < 0.05; ** *P* < 0.01; *** *P* < 0.001; *ns*, not significant). (**E**) Representative immunofluorescence images of YAP and nucleus in hMSCs grown on grown on a 100 kPa gel with constant thickness (larger than 200 μm) in the presence of Cytochalasin at different concentrations stained by YAP antibody (red) and DAPI (blue), respectively. Scale bar: 30 μm. (**F**) The statistical analysis of contact stiffness in hMSCs grown on a 100 kPa gel with different concentrations of Cytochalasin as shown in (E). All error bars are S.E.M. (n = 18, 18, 18, and 15 cells for 0, 0.5, 1, and 2 μM, respectively. *** *P* < 0.001). (**G**) The statistical analysis of YAP N/C ratio in hMSCs grown on a 100 kPa gel with different concentrations of Cytochalasin as shown in (E). All error bars are S.E.M. (n = 18, 18, 18, and 15 cells for 0, 0.5, 1, and 2 μM, respectively. *** *P* < 0.001). (**H**) The statistical analysis of YAP N/C ratio in hMSCs grown on a 100 kPa gel with different contact stiffness as shown in (E). All error bars are S.E.M. (n = 15, 18, 18, and 18 cells for CS = 4.8, 5.3, 6.5, and 10.7 N/m, respectively. ** *P* < 0.01; *** *P* < 0.001). (**I**) Relationship between YAP N/C ratio and contact stiffness or other parameters (ECM stiffness, ECM thickness and cell adhesion area) in hMSCs grown on grown on gradient thickness gels with different stiffness or treated with Cytochalasin. N/C of YAP scales across all the combined datasets as power functions of contact stiffness, with exponent 0.330 ± 0.011 (*r^2^* = 0.789, *P* < 0.001); area, with exponent 0.836 ± 0.047 (*r^2^* = 0.639, *P* < 0.001); thickness, with exponent −0.371 ± 0.022 (*r^2^* = 0.541, *P* < 0.001). Grey area are 95% confidence intervals for the fitted functions. All box-whisker plots show the medians, maxima, minima, upper quartiles and lower quartiles.

Previous studies have reported that YAP activity is regulated by cytoskeletal tension-dependent cell spreading, which leads to variations in the cell adhesion area (*9*). Thus, we further evaluated the use of CS for explaining alterations in YAP activity among cells treated with Cytochalasin, a specific inhibitor of actin polymerization (*29*). There was a dose-dependent decrease in CS and a simultaneous decrease in the YAP N/C ratio among Cytochalasin-treated cells (Fig. 2E–G). The ECM stiffness and thickness were maintained at 100 kPa and ∼200 μm, respectively, for both treated and untreated cells. We therefore inferred that the observed variations in YAP activity resulted from differences in the CS *per se* rather than from differences in ECM stiffness or thickness (Fig. 2H).

We next combined the experimental datasets described above, analyzing data from cells grown on gradient thickness gels with different stiffnesses and data from cells treated with Cytochalasin. Using the combined dataset, we analyzed the fit between the CS and the YAP N/C ratio. In the form of a power function, CS scaled better with the YAP N/C ratio (with an exponent of 0.33 ± 0.011, *r^2^* = 0.789, *P* < 0.001) than other variables did, including ECM stiffness (*r^2^*= 0.057, *P* < 0.001), ECM thickness (*r^2^* = 0.541, *P* < 0.001), and adhesion area (*r^2^* = 0.639, *P* < 0.001) (Fig. 2I). These data demonstrated that the CS could be successfully used to explain YAP mechanosensing in response to different mechanical stimuli.

In addition to the experiments performed in the present study, we also applied CS to interpret YAP mechanosensing in datasets from the literature (*9, 30, 31*). Using published datasets, we found that CS varied significantly between cell types (Supplementary Fig. S2A). However, the YAP N/C ratio consistently scaled well with CS in each cell type (Supplementary Fig. S2B, C). Taken together, these results demonstrated that CS scales better overall with YAP activation than other individual variables in the cell-ECM system do. This is likely because CS integrates the effects of all such variables on cell-ECM interactions into a single value.

### CS could be used to interpret mechanical regulation of stem cell differentiation

The mechanical environment surrounding a cell regulates not only YAP activation but also stem cell differentiation (*4, 6, 32*). Here, we assessed whether CS could be used to interpret mechanical stimulus-induced stem cell differentiation. This was accomplished by analyzing the extent of stem cell differentiation among hMSCs grown on gradient thickness gels. Levels of the osteogenic differentiation markers RUNX2 and Collagen I significantly increased along with increases in CS (Fig. 3A–C). In contrast, levels of the neurogenic markers Nestin and NFL decreased as CS increased, especially when CS was < 1 N/m (Fig. 3D–F). The positive correlations between CS and levels of osteogenic differentiation markers were more pronounced in regions with a high CS (> 10 N/m) than in regions with a low CS (< 1 N/m), whereas the reverse was true of the negative correlations between CS and levels of the neurogenic differentiation markers (Fig. 3, Supplementary Fig. S3). These results demonstrated the sensitivity of stem cell differentiation to variations in CS.

**Fig. 3.**
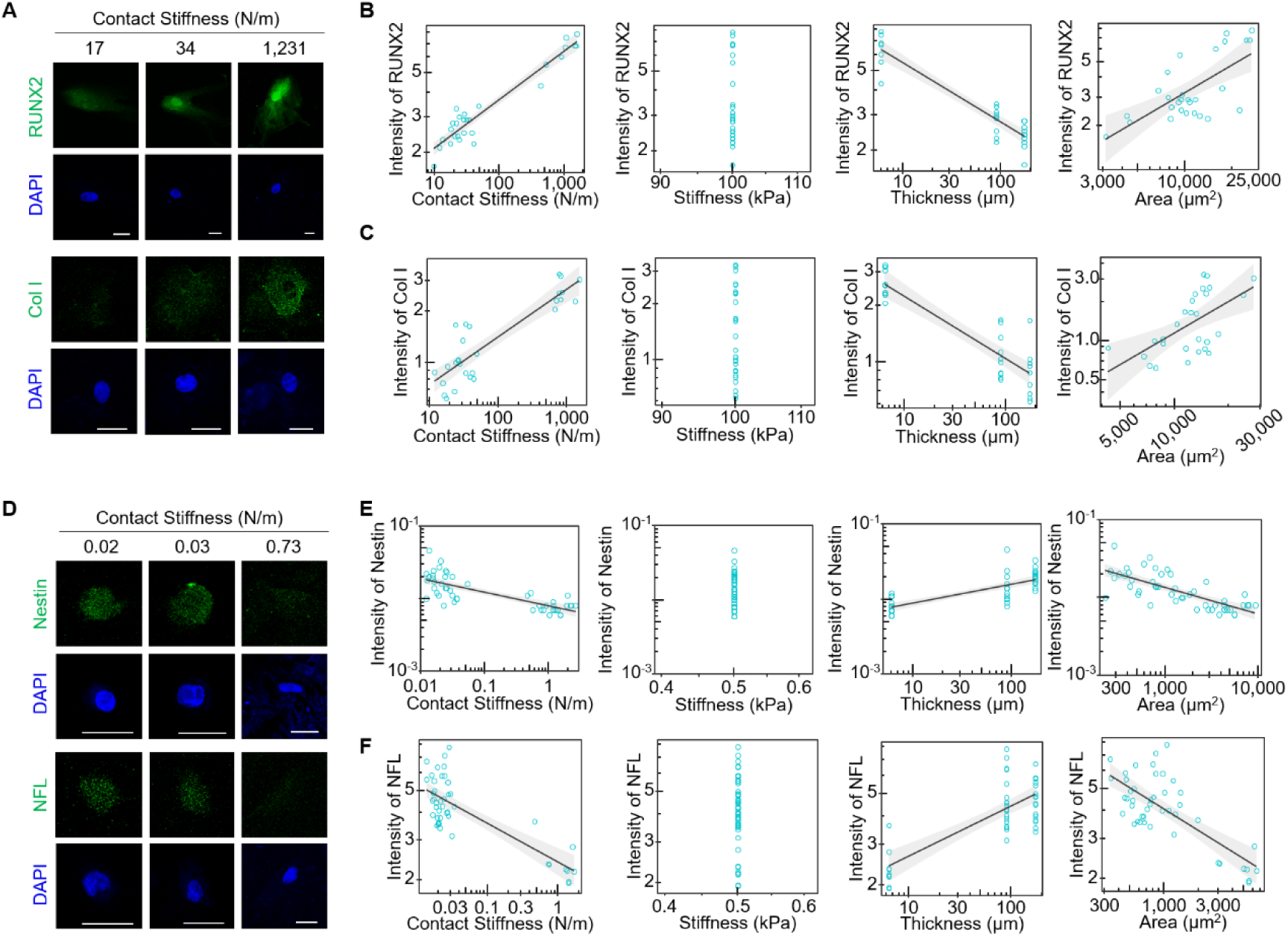
CS could be used to interpret mechanical regulation of stem cell differentiation. (**A**) Representative immunofluorescence images of Runx2 and Collagen I (Col I) in hMSCs grown on a 100 kPa gradient thickness gel for 4 days at different contact stiffness stained by Runx2 or Collagen I antibody (green) and DAPI (blue), respectively. Scale bar: 30 μm. (**B**) and (**C**) Relationship between (B) Runx2 and (C) Collagen I level with contact stiffness or other parameters (ECM stiffness, ECM thickness and cell adhesion area) in hMSCs grown on a 100 kPa gradient thickness gel for 4 days. (B) RUNX2 level scales as power functions of Contact Stiffness, with exponent 0.245 ± 0.011 (*r^2^* = 0.949, *P* < 0.001); Thickness, with exponent −0.297 ± 0.022 (*r^2^* = 0.881, *P* < 0.001); Area, with exponent 0.733 ± 0.140 (*r^2^* = 0.517, *P* < 0.001); Grey area are 95% confidence intervals for the fitted functions. (C) Col I level scales as power functions of Contact Stiffness, with exponent 0.266 ± 0.026 (*r^2^* = 0.828, *P* < 0.001); Thickness, with exponent −0.321 ± 0.033 (*r^2^* = 0.8, *P* < 0.001); Area, with exponent 0.854 ± 0.232 (*r^2^* = 0.539, *P* < 0.001); Grey area are 95% confidence intervals for the fitted functions. (**D**) Representative immunofluorescence images of Nestin and Neurofilament (NFL) and nucleus in hMSCs grown on a 0.5 kPa gradient thickness gel for 4 days at different contact stiffness stained by Nestin or NFL antibody (green) and DAPI (blue), respectively. Scale bar: 30 μm. (**E**) and (**F**) Relationship between (E) Nestin (F) NFL level and contact stiffness or other parameters (ECM stiffness, ECM thickness and cell adhesion area) in hMSCs grown on a 0.5 kPa gradient thickness gel for 4 days. (E) Nestin level scales as power functions of Contact Stiffness, with exponent −0.212 ± 0.044 (*r^2^* = 0.445, *P* < 0.001); Thickness, with exponent 0.278 ± 0.054 (*r^2^* = 0.437, *P* < 0.001); Area, with exponent −0.353 ± 0.064 (*r^2^* = 0.436, *P* < 0.001); Grey area are 95% confidence intervals for the fitted functions. (F) NFL level scales as power functions of Contact Stiffness, with exponent 0.266 ± 0.026 (*r^2^* = 0.462, *P* < 0.001); Thickness, with exponent 0.198 ± 0.045 (*r^2^* = 0.418, *P* < 0.001); Area, with exponent −0.264 ± 0.06 (*r^2^* = 0.388, *P* < 0.001).

We also compared the fit of several variables in the cell-ECM system with the extent of stem cell differentiation. Expression levels of all osteogenic and neurogenic differentiation markers analyzed here scaled better with CS than with variables such as ECM stiffness, ECM thickness, or cell adhesion area (Fig. 3B, C, E, F). These data demonstrated that CS could be successfully used to interpret mechanoregulation of stem cell differentiation. We then explored the usefulness of CS in interpreting the extent of stem cell differentiation induced by treatment with an osteogenic differentiation medium under consistent ECM parameters (*e.g.*, stiffness and thickness). CS varied over time in the presence of the osteogenic differentiation medium during stem cell differentiation (Supplementary Fig. S4A–C). Importantly, RUNX2 levels increased along with the CS, and the CS scaled well with RUNX2 expression (Supplementary Fig. S4D–F). ECM stiffness and thickness were both maintained at a steady state over time while the CS changed; differences in the CS alone, rather than in ECM stiffness or thickness, therefore accounted for stem cell differentiation. These results indicate that the use of CS provides an opportunity to reassess what is known about the mechanisms underlying the effects of ECM mechanics on stem cell differentiation.

### A CS-based motor clutch model for investigating the cross-scale ECM mechanosensing

Previous studies have demonstrated that cell architecture is regulated by the activity of adhesion molecules (*e.g.*, FA plaques) and the cytoskeleton (*e.g.*, actin filaments, F-actin) (*33, 34*). The mechanosensing mechanisms by which cells respond to ECM mechanics have previously been explained with molecular motor clutch models (*21, 35*), in which local ECM stiffness is perceived through the assembly of FA plaques and balanced by pulling forces actuated by F-actin (Fig. 4A). In these models, the local reaction force of the ECM is related to local ECM deformation as a function of stiffness. However, contact mechanics theory (*22*) indicates that local ECM deformation depends on both the local force (*i.e.*, the force exerted by local FAs) and the non-local force (from non-local FAs in the cell). This indicates that a cross-scale model would be necessary to interpret the mechanism by which cell-ECM interactions influence the assembly of sub-cellular elements such as FA plaques and F-actin.

**Fig. 4.**
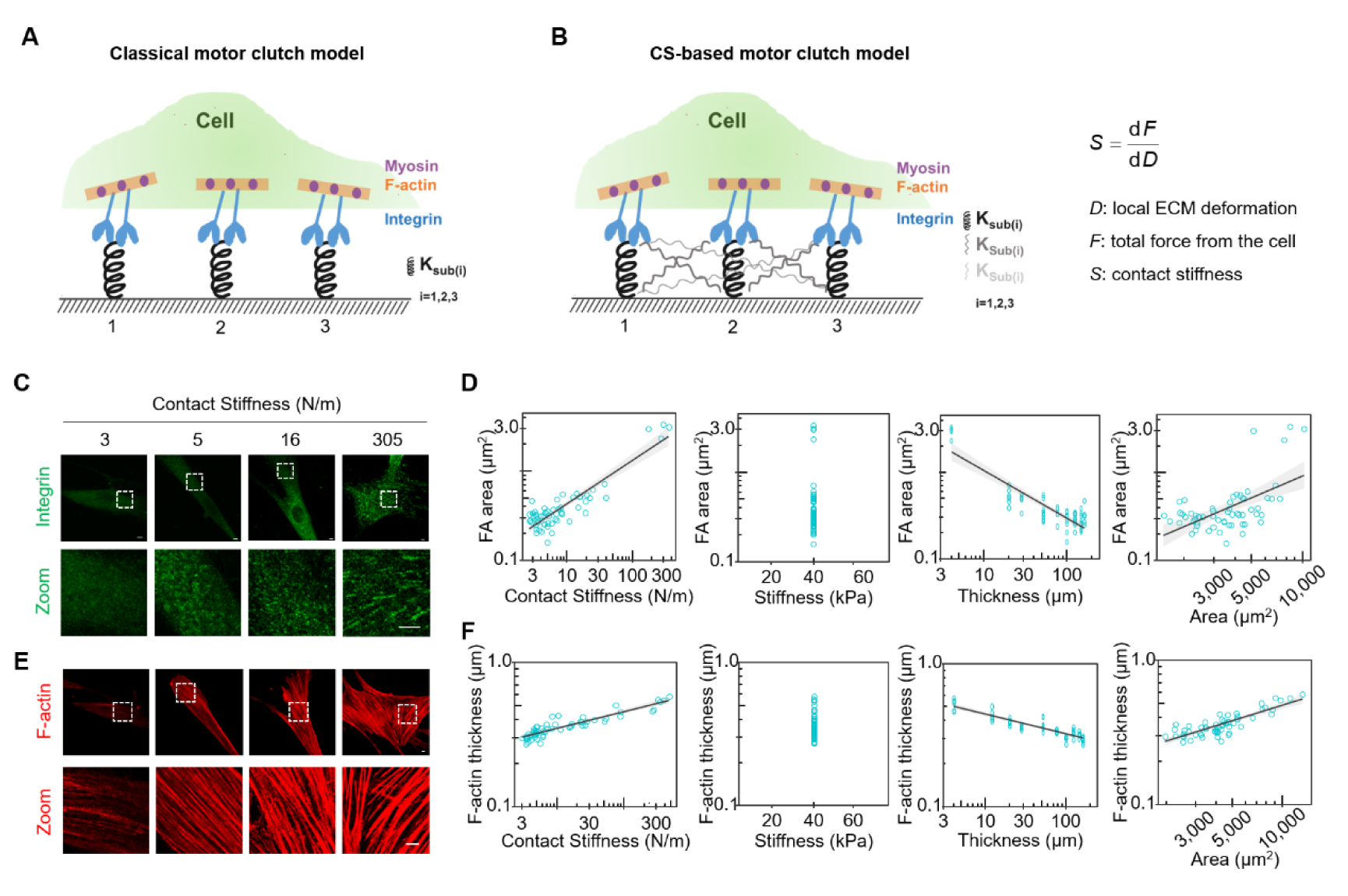
A CS-based motor clutch model for investigating the cross-scale ECM mechanosensing. (**A**) Schematic diagram of cell-ECM interactions illustrated by a classical local motor clutch model. (**B**) Schematic diagram of cell-ECM interactions illustrated by a contact stiffness-based non-Local motor clutch model. The variation in ECM local deformation (*D*) is related to the change of the total force (*F*) from the cell by the CS (*S*). (**C**) Representative images of Integrin in hMSCs grown on a 40 kPa gradient thickness gel at different contact stiffness stained by β1 Integrin antibody. The fluorescence images below are enlarged views of the top white dotted boxes. Scale bar: 5 μm. (**D**) Relationship between focal adhesion area (analyzed by β1 integrin antibody staining) and contact stiffness or other parameters (ECM stiffness, ECM thickness and cell adhesion area) in hMSCs grown on a 40 kPa gradient thickness gel. FA area scales as power functions of Contact Stiffness, with exponent 0.582 ± 0.021 (*r^2^*= 0.934, *P* < 0.001); Thickness, with exponent −0.819 ± 0.031 (*r^2^* = 0.934, *P* < 0.001); Area, with exponent 2.295 ± 0.256 (*r^2^* = 0.554, *P* < 0.001); Grey area are 95% confidence intervals for the fitted functions. (**E**) Representative images of F-actin in hMSCs grown on a 40 kPa gradient thickness gel at different contact stiffness stained by Phalloidin. The fluorescence images below are an enlarged view of the top white dotted boxes. Scale bar: 5 μm. (**F**) Relationship between the extent of F-actin assembly and contact stiffness or other parameters (ECM stiffness, ECM thickness and cell adhesion area) in hMSCs grown on a 40 kPa gradient thickness gel. F-actin thickness scales as power functions of Contact Stiffness, with exponent 0.115 ± 0.006 (*r^2^* = 0.815, *P* < 0.001); Thickness, with exponent −0.143 ± 0.008 (*r^2^* = 0.828, *P* < 0.001); Area, with exponent 0.396 ± 0.027 (*r^2^* = 0.762, *P* < 0.001); Grey area are 95% confidence intervals for the fitted functions.

As discussed above, CS (S) defines the relationship between local ECM deformation (D) and the total force from the cell (F, which is composed of local and non-local forces and equals the reaction force of the ECM on the cell) as *S* = d *F* / d *D*. Because of this relationship and our finding that CS scaled with both YAP activity and the extent of stem cell differentiation, we here propose a CS-based model with reference to classical motor clutch models (Fig. 4B). This model suggests that variations in local ECM deformation arise from the total force imposed by the cell (including local and non-local forces), which is dependent on CS. The reaction force provided by the local ECM causes force-dependent assembly of FA plaques and actin filaments (*2*). This may lead to alterations in cell architecture (*e.g.*, adhesion area or shape) by mechanotransduction, further affecting the whole-cell force. Because CS is determined by the cell adhesion area and shape in addition to ECM stiffness and thickness, variations in cell architecture change the CS in turn, ultimately affecting the relationship between local ECM deformation and the total force from the cell. Therefore, a cross-scale regulatory loop between local ECM deformation and the total force between the cell and the ECM is established during cell-ECM interactions (Fig. 4B).

To verify this CS-based motor clutch model, we assessed the relationships between CS and assemblies of FA plaques or F-actin. Immunofluorescent staining showed that FA plaque size and F-actin thickness both increased along with CS in hMSCs cultured on gradient thickness gels (Fig. 4C, E). In combined datasets, we found that CS scaled better in the form of power functions with FA plaque size (*r^2^*= 0.934, *P* < 0.001) and F-actin assemblies (*r^2^* = 0.86, *P* < 0.001) than other individual variables did (*e.g.*, ECM stiffness or thickness; cell adhesion area or shape) (Figure 4d, f). These results indicate that the CS-based motor clutch model enables interpretation of the effects of ECM mechanics on dynamic FA plaque and cytoskeleton assembly during cell-ECM interactions. Thus, the CS-based model could likely be used as a frame of reference to further explore ECM mechanosensing.

### The CS-based motor clutch model reveals the contributions of time-dependent cell architecture evolution to stem cell differentiation

Two remarkable features of stem cell differentiation are long-lasting changes in gene expression and time-dependent evolutions in cellular architecture (*36*). However, the mechanism by which cell architecture evolves over time and its contributions to the progress of stem cell differentiation remain unclear. Our CS-based motor clutch model suggested a relationship between cell architecture and the reaction force from the ECM. Thus, we sought to delineate the involvement of cell architecture in stem cell differentiation based on CS. We first compared the adhesion areas of hMSCs grown on ECMs of different thicknesses at distinct time points. Overall, there were significant increases in adhesion area over time (Fig. 5A, B), which led to clear time-dependent increases in CS (Fig. 5C). Similar variations in the adhesion area and CS over time were also observed among hMSCs grown on gels with consistent ECM mechanics that were treated with osteogenic differentiation medium (Supplementary Fig. S4A–C). These results indicated that the evolution of cell architecture led to a corresponding change in CS over time during stem cell differentiation.

**Fig. 5.**
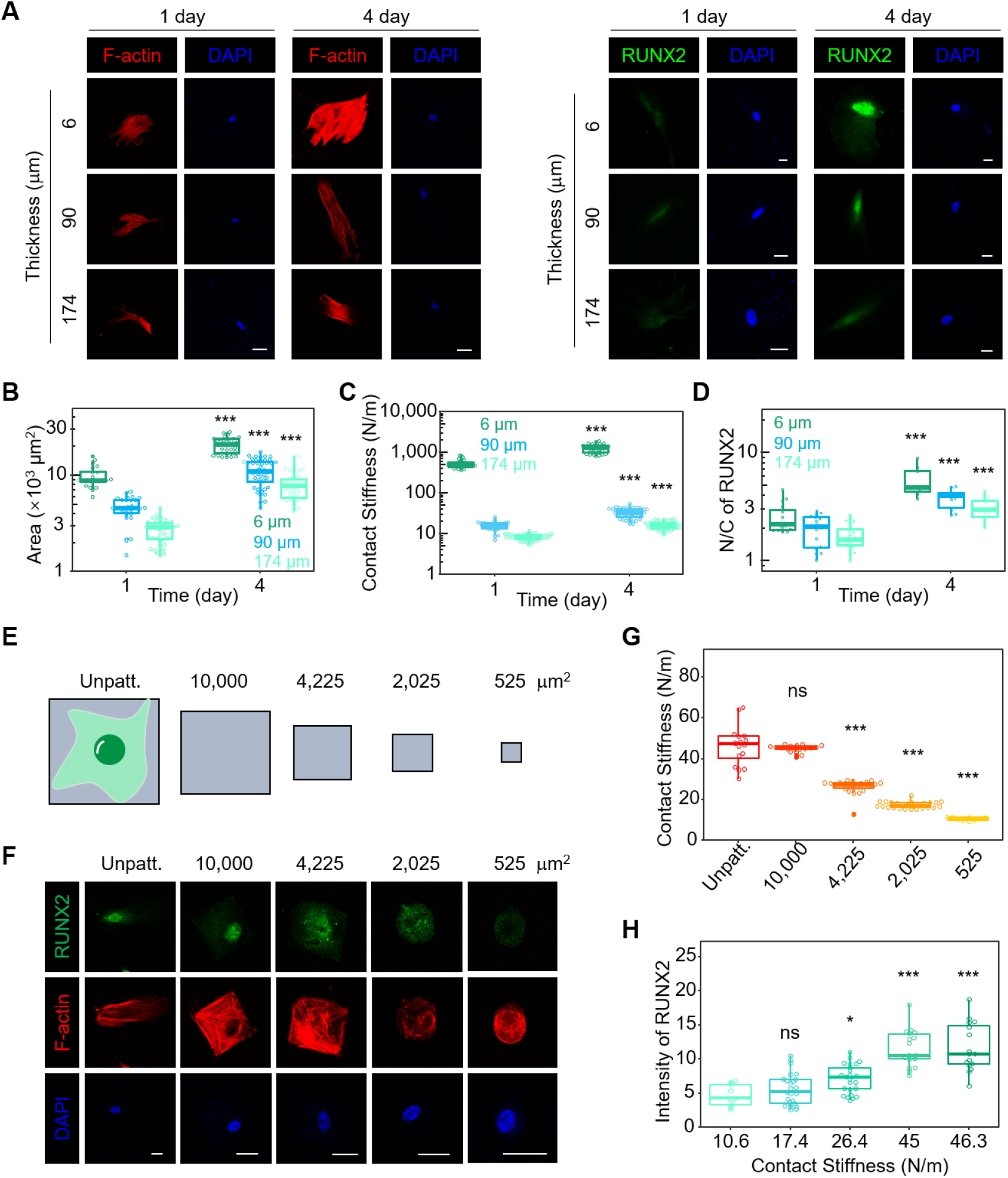
The CS-based motor clutch model reveals the contributions of time-dependent cell architecture evolution to stem cell differentiation. (**A**) Representative images of F-actin or RUNX2 and nucleus in hMSCs grown on a 100 kPa gradient thickness gel at distinct thickness regions for 1 or 4 days stained by phalloidin (red) or RUNX2 (green) and DAPI (blue), respectively. Scale bar: 30 μm. (**B**) The statistical analysis of cell adhesion area in hMSCs grown on a 100 kPa gradient thickness gel at distinct thickness regions for 1 or 4 days as shown in (A). All error bars are S.E.M. (1 day: n = 21 cells for 6 μm; n = 28 cells for 90 μm; n = 32 cells for 174 μm. 4 day: n = 33 cells for 6 μm; n = 46 cells for 90 μm; n = 42 cells for 174 μm. *** *P* < 0.001). (**C**) The statistical analysis of contact stiffness in hMSCs grown on a 100 kPa gradient thickness gel at distinct thickness regions for 1 or 4 days as shown in (A). All error bars are S.E.M. (1 day: n = 21 cells for 6 μm; n = 28 cells for 90 μm; n = 32 cells for 174 μm. 4 day: n = 33 cells for 6 μm; n = 46 cells for 90 μm; n = 42 cells for 174 μm. *** *P* < 0.001). (**D**) The statistical analysis of RUNX2 N/C ratio in hMSCs grown on a 100 kPa gradient thickness gel at different contact stiffness for 1 or 4 days as shown in (A). All error bars are S.E.M. (1 day: n = 20 cells for 6 μm; n = 16 cells for 90 μm; n = 22 cells for 174 μm. 4 day: n = 8 cells for 6 μm; n = 10 cells for 90 μm; n = 11 cells for 174 μm. *** *P* < 0.001). (**E**) Grey patterns show the relative size of micropatterned fibronectin islands on which cells were grown. Outline of a cell is shown superimposed to the leftmost unpatterned area (Unpatt.). (**F**) Representative images of RUNX2 (green), F-actin (red) and nucleus (blue) in hMSCs grown on micropatterned fibronectin islands. Scale bar: 30 μm. (**G**) The statistical analysis of contact stiffness in hMSCs grown on a micropatterned fibronectin islands as shown in (f). All error bars are S.E.M. (Unpatt.: n = 16 cells; 10000 μm^2^: n = 18 cells; 4225 μm^2^: n = 25 cells; 2025 μm^2^: n = 27 cells; 525 μm^2^: n = 9 cells. *** *P* < 0.001; *ns*, not significant; compared to Unpatt.). (**H**) The statistical analysis of intensity of RUNX2 in hMSCs grown on a micropatterned fibronectin islands as shown in (F). All error bars are S.E.M. (n = 9, 27, 25, 18, and 16 cells for CS = 10.6, 17.4, 26.4, 45, and 46.3 N/m, respectively. *** *P* < 0.001; *ns*, not significant; compared to Unpatt.).

Based on the CS-based motor clutch model, the increased CS during osteogenic differentiation of hMSCs indicated that cells were subject to a greater reaction force over time from the same local ECM deformation, which may have affected the progress of differentiation. To evaluate this hypothesis, we analyzed expression levels of osteogenic differentiation markers. Immunofluorescent staining showed that RUNX2 levels were significantly increased over time among all hMSCs, regardless of ECM thickness. Increases in RUNX2 levels were greater among cells grown on a thin ECM than on a thick ECM (Fig. 5A, D). Similar results were observed in hMSCs treated with osteogenic differentiation medium under constant ECM mechanics (Supplementary Fig. S4E, F). We then inhibited time-dependent CS increases by culturing hMSCs on micropatterned islands with different adhesion areas (Fig. 5E). Immunofluorescent staining showed a consistent decrease in both CS and RUNX2 levels in the physically-constrained cells (Fig. 5F–H), indicating that the evolution of CS over time markedly affected stem cell differentiation. Thus, the evolution of cell architecture over time appeared to alter the CS and the reaction force provided to the cell by the ECM during cell-ECM interactions. These factors combined to strongly affect stem cell differentiation.

### The CS-based motor clutch model predicts the effects of a non-adjacent layer of composite ECM on cellular mechanosensing

The ECMs of living organisms are not homogeneous materials. ECMs may form layered composites with hierarchically-structured assemblies of diverse biological materials (*12, 37*). Depending on the topographical positions, ECMs are generally divided into a pericellular matrix (*e.g.*, basement membranes [BMs]) and an underlying interstitial matrix (*e.g.*, ECMs in connective tissues). The pericellular matrix is in direct contact with cells and provides an adhesive microenvironment for resident cells (*e.g*., epithelial cells), whereas the interstitial matrix underlies the pericellular matrix and supports mechanical tissue integrity (*37*). Previous studies of mechanical-based cell behavioral regulation by ECMs have focused on the ECM layer located adjacent to the cell (*e.g.,* the pericellular matrix, here referred to as the adjacent ECM layer). However, the CS-based motor clutch model indicated that non-adjacent ECM layers (such as the interstitial matrix) would also affect CS of the composite ECM (*38, 39*), influencing the reaction force of the ECM on the cell when the ECM was deformed.

To experimentally validate these predictions, we prepared a two-layered ECM with a thickness gradient. This consisted of an adjacent ECM layer with a constant stiffness of 10 kPa and a non-adjacent ECM layer with either 40 kPa stiffness (the 10/40-kPa gel) or 100 kPa stiffness (the 10/100-kPa gel) (Fig. 6A). Using this experimental system, we cultured hMSCs and analyzed the responses of cells to variations in non-adjacent ECM layer mechanics. Compared with cells cultured on the 10/40-kPa gel, cells cultured on the 10/100-kPa gel exhibited larger cell adhesion areas at distinct regions of thickness (Fig. 6B, C). We then calculated CS based on measurements of individual variables. Notably, CS was greater on the 10/100-kPa gel than on the 10/40-kPa gel at distinct thickness regions (Fig. 6D). These results showed that CS varied with the stiffness of the non-adjacent ECM layer.

**Fig. 6.**
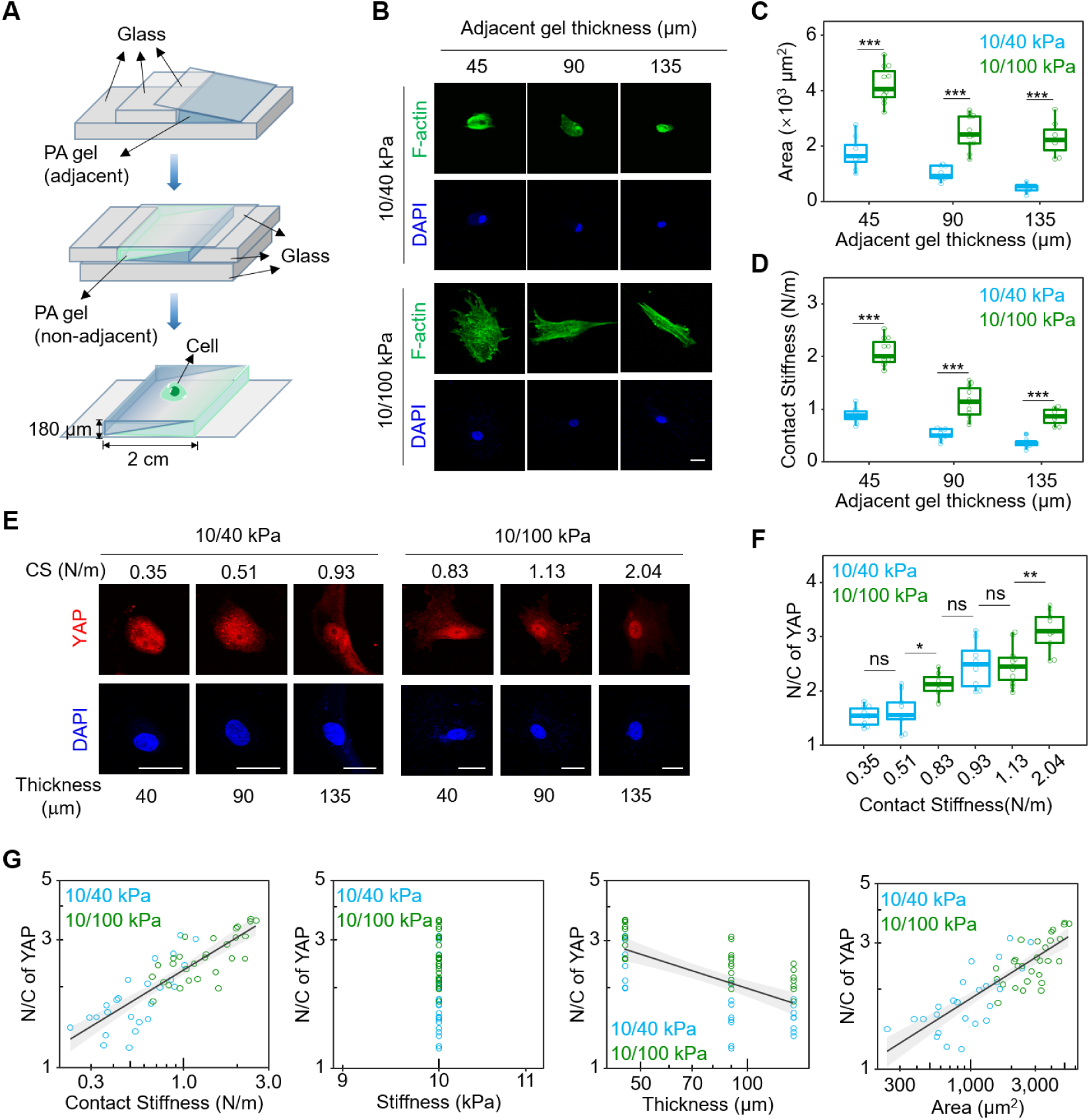
The CS-based motor clutch model predicts the effects of a non-adjacent layer of composite ECM on cellular mechanosensing. (**A**) Schematic diagram of the fabrication of a two-layered ECM with gradient thickness composed of a cell adjacent layer and a cell non-adjacent layer. (**B**) Representative images of F-actin and nucleus in hMSCs grown on two-layered gradient thickness ECMs composed of a cell adjacent layer with constant Young’s modulus of 10 kPa and a non-adjacent layer with varied Young’s modulus of 40 (10/40 kPa) or 100 kPa (10/100 kPa) gel at distinct thickness regions stained by phalloidin (green) and DAPI (blue), respectively. Scale bar: 30 μm. (**C**) The statistical analysis of cell adhesion area in hMSCs grown on two-layered gradient thickness ECM at distinct thickness regions as shown in (B). All error bars are S.E.M. (n = 8 and 11 cells for 45 μm thickness of 10/40 kPa and 10/100 kPa, respectively; n = 9 and 11 cells for 90 μm thickness of 10/40 kPa and 10/100 kPa, respectively; n = 8 and 7 cells for 175 μm thickness of 10/40 kPa and 10/100 kPa, respectively. *** *P* < 0.001). (**D**) The statistical analysis of contact stiffness in hMSCs grown on two-layered gradient thickness ECMs at distinct thickness regions as shown in (B). All error bars are S.E.M. (n = 8 and 11 cells for 45 μm thickness of 10/40 kPa and 10/100 kPa, respectively; n = 9 and 11 cells for 90 μm thickness of 10/40 kPa and 10/100 kPa, respectively; n = 8 and 7 cells for 175 μm thickness of 10/40 kPa and 10/100 kPa, respectively. *** *P* < 0.001). (**E**) Representative immunofluorescence images of YAP and nucleus in hMSCs grown on two-layered gradient thickness ECMs at different contact stiffness stained by YAP antibody (red) and DAPI (blue), respectively. Scale bar: 30 μm. (**F**) The statistical analysis of YAP N/C ratio in hMSCs grown on two-layered gradient thickness ECMs at different contact stiffness as shown in (E). All error bars are S.E.M. (n = 10, 9, 7, 8, 11, and 9 cells for CS = 0.35, 0.51, 0.83, 0.93, 1. 13, and 2.04 N/m, respectively. *** *P* < 0.001; *ns*, not significant; compared to Unpatt.). * *P* < 0.05; ** *P* < 0.01; *** *P* < 0.001; *ns*, not significant). (**G**) Relationship between YAP N/C ratio and contact stiffness or other parameters (ECM stiffness, ECM thickness and cell adhesion area) in hMSCs grown on two-layered gradient thickness ECMs. All box-whisker plots show the medians, maxima, minima, upper quartiles and lower quartiles. N/C of YAP scales across all the datasets as power functions of Contact Stiffness, with exponent 0.395 ± 0.035 (*r^2^* = 0.72, *P* < 0.001); Thickness, with exponent −0.417 ± 0.065 (*r^2^* = 0.446, *P* < 0.001); Area, with exponent 0.342 ± 0.034 (*r^2^* = 0.687, *P* < 0.001); Grey area are 95% confidence intervals for the fitted functions.

To investigate whether cells could sense mechanical stimuli from the non-adjacent ECM layer, we analyzed YAP activity among hMSCs cultured on the two-layered ECMs. Immunofluorescent staining showed that cells cultured on the 10/100-kPa gel had significantly higher YAP N/C ratios than cells cultured on the 10/40-kPa gel at distinct thickness regions (Fig. 6E, F). Interestingly, when comparing the datasets from the 10/40-kPa and 10/100-kPa gels at different thicknesses, we found that the YAP N/C ratios were similar only if the CS values were comparable, despite differences in the ECM thickness or non-adjacent layer stiffness. For example, the YAP N/C ratios were similar between cells grown on a 10/40-kPa gel that was 6 μm thick (CS = 0.93 N/m) and those grown on a 10/100-kPa gel that was 40 μm thick (CS = 0.83 N/m) (Fig. 6E, f). Moreover, the YAP N/C ratio scaled better with CS than with other individual variables (namely ECM stiffness or thickness or cell adhesion area) (Fig. 6G). These findings reveal that the regulatory effects of mechanical stimuli from a non-adjacent ECM layer on cells could be interpreted as influencing the reaction force provided by the ECM to the cell due to its contribution to CS.

## DISCUSSION

In cell-ECM interactions, cells are continuously exposed to physical forces imposed by the ECM. Understanding the mechanisms by which such physical forces are generated requires an understanding of the relationship between ECM deformation and the ECM reaction force on a cell (*1–3*). CS defines the relationship between the local ECM deformation and the force from the cell (which equals the reaction force of the ECM on a living cell). We therefore proposed a CS-based frame of reference to investigate ECM mechanosensing. Our experiments reveal the advantages of modeling such dynamics with CS rather than individual variables (*e.g.*, ECM stiffness, ECM thickness, or cell adhesion area) to interpret ECM mechanosensing. The experimental data demonstrated that YAP activation and the extent of stem cell differentiation triggered by tuning these individual variables can ultimately be understood as modulations of CS (Supplementary Fig. S5). Thus, using CS as a frame of reference could provide opportunities for understanding the common mechanisms underlying the mechanoregulation of multiple ECM stimuli.

Based on the multi-scale features through which cells interact with the ECM, we proposed a CS-based model built on classical motor clutch models; this showed that local ECM deformation generated a reaction force not only on the FAs contacting a local position, but also on the entire cell in a long-range, non-local manner. Such a force-deformation relationship, according to contact mechanics theory (*22*), can be defined by CS, which integrates the effects of ECM stiffness and thickness and the cell adhesion area and shape on cell-ECM interactions at the cellular level. The effects of ECM prestress on cellular behaviors have been addressed in previous studies (*40*), in which the strain energy of the ECM has been suggested as the mechanical variable that is sensed by a cell. We used the CS-based model presented here to interpret the effects of ECM prestress on cells. In the presence of prestress, the CS can be expressed as 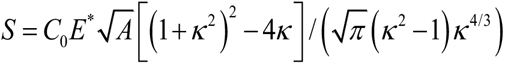 where *κ* ≥ 1 measures the extent of pre-tension (*38*) (Supplementary Fig. S5). This analytical solution shows that pre-tension increases the CS and leads to a larger adhesion area, consistent with previous reports (*40*).

At a larger scale, in the context of collective cell behaviors (such as those of cell sheets) (*11, 13*), our CS-based motor clutch model implies that not only the variables mentioned above, but also the in-plane dimension of a cell sheet, will affect interactions of the cell sheet with the ECM. For example, for a multi-layered ECM, our model predicted that significant effects on collective cell behavior would arise when the layer had a depth less than or equal to the in-plane cell sheet dimensions. This interesting prediction will require further validation.

The CS-based model predicted that dynamic variations occurred in parameters from mechanosensitive assembly of single FA plaques to the overall cell architecture, including the cell adhesion area and shape over time. All of these factors influenced the CS and therefore ultimately influenced interactions between cells and the ECM. Therefore, the CS-based model may help to explain the dynamic features involved in ECM mechanosensing, mechanotransduction, and mechanoresponse. Our experiments showed that variations in the CS range differed significantly between different types of cells grown on the same gradient thickness gel. hMSCs had the widest CS range of all the cell types analyzed (Supplementary Fig. S6). This indicated that time-dependent architectural reorganization was more active in stem cells than in other differentiated cells in response to mechanical stimuli, which contributed to their differentiation. Notably, time-dependent variations in CS may have also stemmed from time-dependent ECM deformations (*41*); in this sense, the CS-based model can therefore be used to interpret the effects of ECM viscoelasticity on cells.

Taken together, our findings demonstrated that CS is a key mechanical variable that can be sensed and actively tuned by cells. This variable provides a frame of reference for future exploration of cross-scale and dynamic mechanisms of mechanical cell-ECM interactions and the effects of various mechanical stimuli in ECMs. This framework will enable a deeper, more comprehensive understanding of the mechanosensing mechanisms that respond to the ECM as part of many physiological and pathological processes, including embryonic development, tissue regeneration, and tumor metastasis.

## ACKNOWLEDGEMENT

We thank Bin Chen (Zhejiang University), Jiliang Hu (Tsinghua University), and Bo Wang (Chinese Academy of Sciences) for discussions during the preparation of this paper. This study was supported by the National Natural Science Foundation of China (NSFC) (11972206, 12222201, 82273500, 11572179, 11432008), the National Key R&D Program of China (2017YFA0506500) and Fundamental Research Funds for the Central Universities (ZG140S1971).

## AUTHOR CONTRIBUTIONS

Y.P.C. suggested the CS-based model, J.D. and Y.P.C. designed and supervised the experiments, P.Z., Y.Z., Y.X.J and L.G. performed the experiments, Z.Y.Z. and M.W.J. developed the finite element model. Y.N.G. and J.L.C. helped with the data analysis and all the authors took part in the data analysis. J.D. and Y.P.C. wrote the paper.

## METHODS

### Fabrication of PA hydrogels with gradient thickness

Single layer PA hydrogel molds were fabricated with two glass slides (70 × 24 × 1 mm and 24 × 24 × 0.18 mm, CITOTEST) as shown in Figure 1b. Double layer hydrogel molds were fabricated with one 70 × 24 × 1 mm glass slide and two 24 × 24 × 0.18 mm as shown in Figure 6a. PA hydrogel preparation was performed according to previous studies (*42*). The mixed liquid cover glass liquid was added to the mold. After polymerization, single layer PA hydrogel was removed from the mold and placed into a culture dish and stored at 4 °C. Double layer PA hydrogels was fabricated on a single-layer PA hydrogel as the mold. After polymerization, the layered gel was removed from the mold and placed into a culture dish and stored at 4 °C. The prepared PA gels were functionalized according to the method described previously (*42*).

### Cell Culture

Human MSCs, HeLa cells, MDCK cells and MEFs were cultured in Dulbecco’s modified eagle medium (DMEM) (containing with 4.5 g L^−1^ glucose, l-glutamine, and sodium pyruvate) supplemented with 10% fetal bovine serum (FBS; Life Technologies, CA, USA), 100 IU mg^−1^ penicillin–streptomycin (Life Technologies, CA, USA), and 1% (v/v) nonessential amino acids (NEAA; Life Technologies, CA, USA) at 37°C and 5% CO_2_.

### Immunocytochemical Staining

Cells were fixed with 4% paraformaldehyde for 30 min at room temperature. After cell membrane was broken using 0.2% Triton X-100 for 10 min at room temperature, samples were blocked with 5% bovine serum albumin for 2 h. The samples were then incubated with primary antibodies against β1 integrin (1:200; Abcam), YAP (1:200; Abcam), RUNX2 (1:200; Abcam), Collagen I (1:200; Abcam), Nestin (1:200; Abcam), and NFL (1:200; Abcam) at 4 °C overnight. The samples were then incubated with secondary antibodies with fluorescence FITC (1:500; Abcam), TRITC (1:500; Abcam) or cy5 (1:200; Abcam) for 2 h at 37 °C. After staining cell nuclei and F-actin by DAPI (1:1000; Sigma) and phalloidin respectively samples were viewed under a Leica SP8 confocal microscopy system. Quantification of the immunofluorescence signal was performed by ImageJ software.

### Statistical Analysis

The data analysis was processed by one-way analysis of variance using Excel and SPSS. Data were presented as mean ± standard error of mean (S.E.M.), and boxplot format data were presented as median ± min/max, as indicated in the corresponding figure legends. Sample size (n) for each statistical analysis was presented in the corresponding figure legends. Statistical significance was determined by the two-tailed Student’s t-test and one-way ANOVA with Tukey’s correction, as indicated in the corresponding figure legends. Tukey’s post-hoc test was used for multiple post-hoc comparisons to determine the significance between the groups after one-way analysis of variance (ANOVA). *P* < 0.05 was considered statistically significant. Statistical analysis was carried out using commercial software IBM SPSS Statistics 22.

### Calculation of contact stiffness

In cases where the ECM is far thicker than a cell (> 10-fold), CS was calculated with 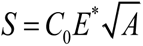, where *C*_0_ is a constant and taken as 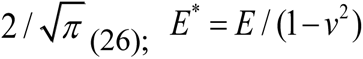(26); *E*\* = *E* / (1− *v*^2^) is the plane-strain modulus (which defines ECM stiffness), *E*\* is measured with indentation tests (Optics11, PIUMA) and tip with radius of 10*μm* is used.; and *A* is the cell adhesion area measured in experiments. When the ECM thickness must be considered (i.e., ECM thickness is on the same order as the dimensions of the cell), CS was calculated with 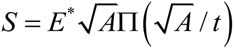, where Π is a dimensionless function as determined by Cao *et al*. (*43*) and *t* is the ECM thickness. To explore the effect of non-adjacent layer on cell behavior, a two-layer hydrogel resting on the stiff substrate has been used. To calculate the contact stiffness of this two-layer ECM, for measured contact area *A*, layer thickness *t*_1_,*t*_2_, elastic moduli *E*_1_,*E*_2_, finite element simulations as described below were performed to determine the contact stiffness.

### Finite element simulations

To determine the contact stiffness and the effect of contact geometry on the contact stiffness, the Finite Element analyses (FEA) was performed using Abaqus/standard (Abaqus 6.14, Dassault Systèmes®). In the simulation, a three-dimensional finite element model has been built, for a given contact area, we imposed normal displacement in the contact area and calculated the reaction force of the two-layer ECM, based on which the contact stiffness was calculated. We used a uniform mesh grid (element size 0.1 mm) in the contact area, and gradient grid in other areas (element size from 0.1mm to 1.0 mm). The hybrid element-C3D10MH (10-node modified tetrahedron element with hourglass control) was used in the whole model.

To study the effect of contact geometry on the contact stiffness, we changed the ratio of a/b of the elliptical contact region (as shown in Fig. S7A), where a and b are the semi-major and semi-minor axis, respectively. The ECM was assumed to be nearly incompressible (Poisson’s ratio is taken as 0.49995) and Young’s modulus is taken as 10 kPa. The displacement boundary was set in the contact area with lower boundary fixed. From the given displacement and calculated reaction force, the contact stiffness was calculated.

## FIGURES

**Fig. S1.**
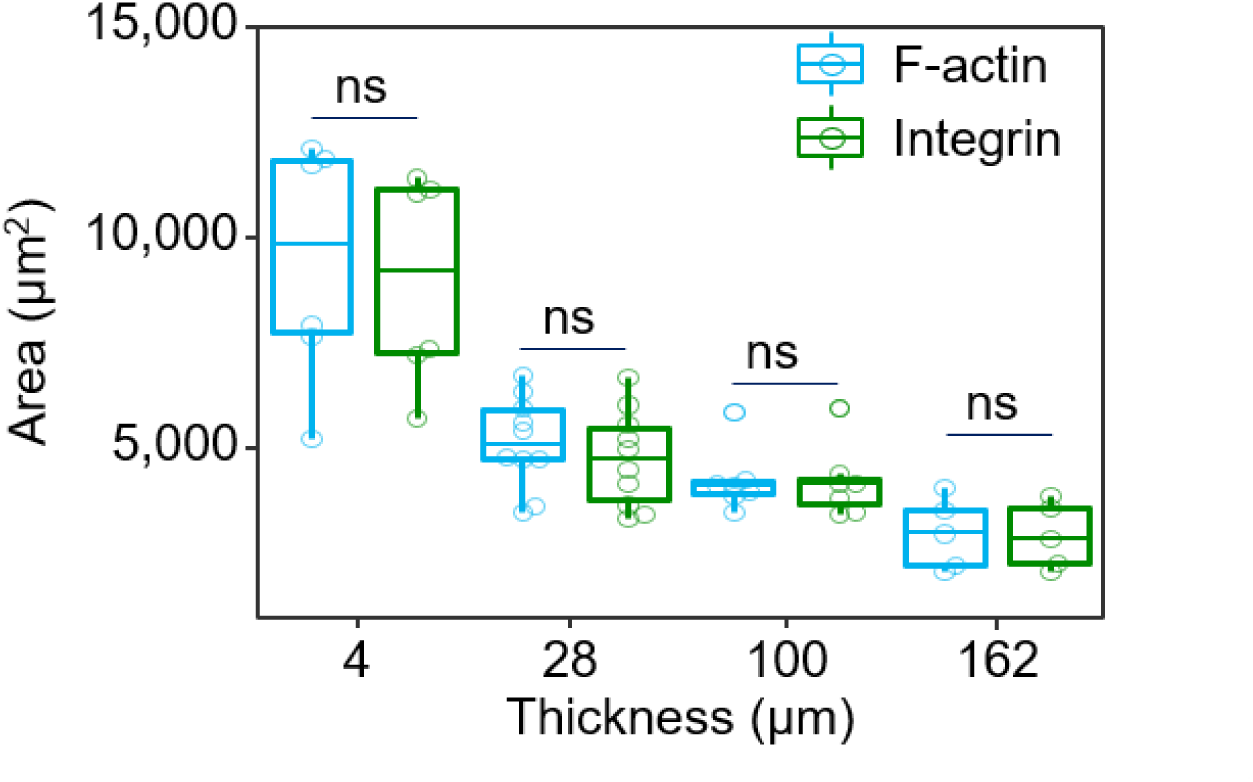
Consistent results of cell adhesion area measured by F-actin and integrin staining. The statistical analysis of cell adhesion area measured by F-actin (labeled by phalloidin) and integrin (labeled by β1 integrin antibody) staining in hMSCs grown on 40 kPa gradient thickness gels for 1 days. All error bars are S.E.M. (F-actin: n = 6, 10, 7, and 5 cells for 4, 28, 100, and 162 μm thickness, respectively; Integrin: n = 6, 10, 7, and 5 cells for 4, 28, 100, and 162 μm thickness, respectively. *ns*, not significant).

**Fig. S2.**
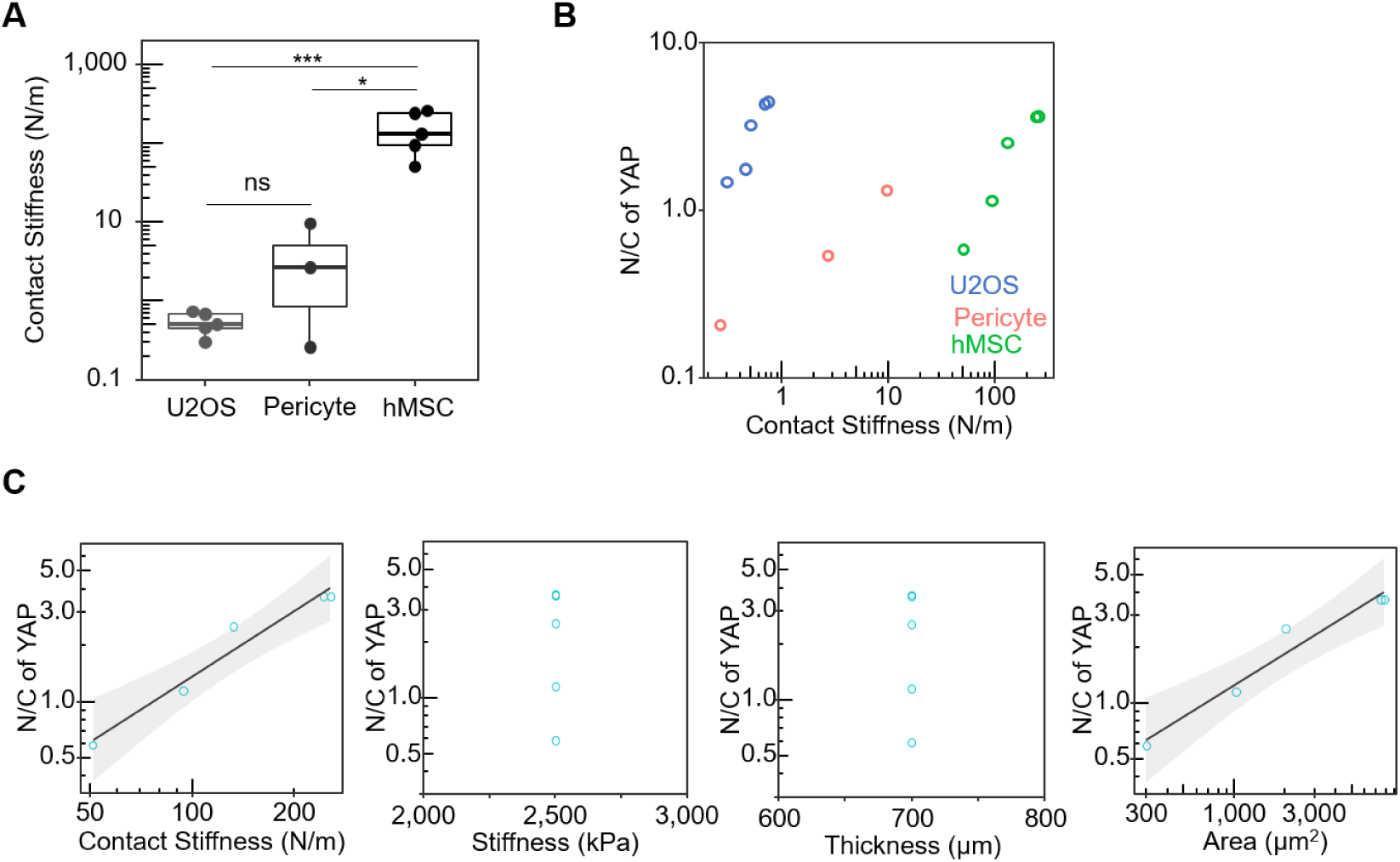
Contact stiffness scales with YAP activity from datasets in the literature. (**A**) The statistical analysis of CS in U2OS cells (*31*), Pericytes (*30*) and hMSCs (*9*) grown on hydrogel or PDMS gel for 1 day. All error bars are S.E.M. (U2OS: n = 5 cells; Pericyte: n = 3 cells; hMSC: n = 5 cells) * *P* < 0.05; *** *P* < 0.001; *ns*, not significant. (**B**) The statistical analysis of YAP N/C ratio in U2OS cells, Pericytes and hMSCs grown on hydrogel or PDMS gel for 1 day. (**C**) Relationship between YAP N/C ratio and contact stiffness or other parameters (ECM stiffness, ECM thickness and cell adhesion area) in in hMSCs grown on PDMS gel for 1 day. N/C of YAP in hMSCs scales as power functions of Contact Stiffness, with exponent 0.967 ± 0.181 (*r^2^* = 0.945, *P* < 0.001); Area, with exponent 0.47 ± 0.089 (*r^2^* = 0.945, *P* < 0.001); Grey area are 95% confidence intervals for the fitted functions. All box-whisker plots show the medians, maxima, minima, upper quartiles and lower quartiles.

**Fig. S3.**
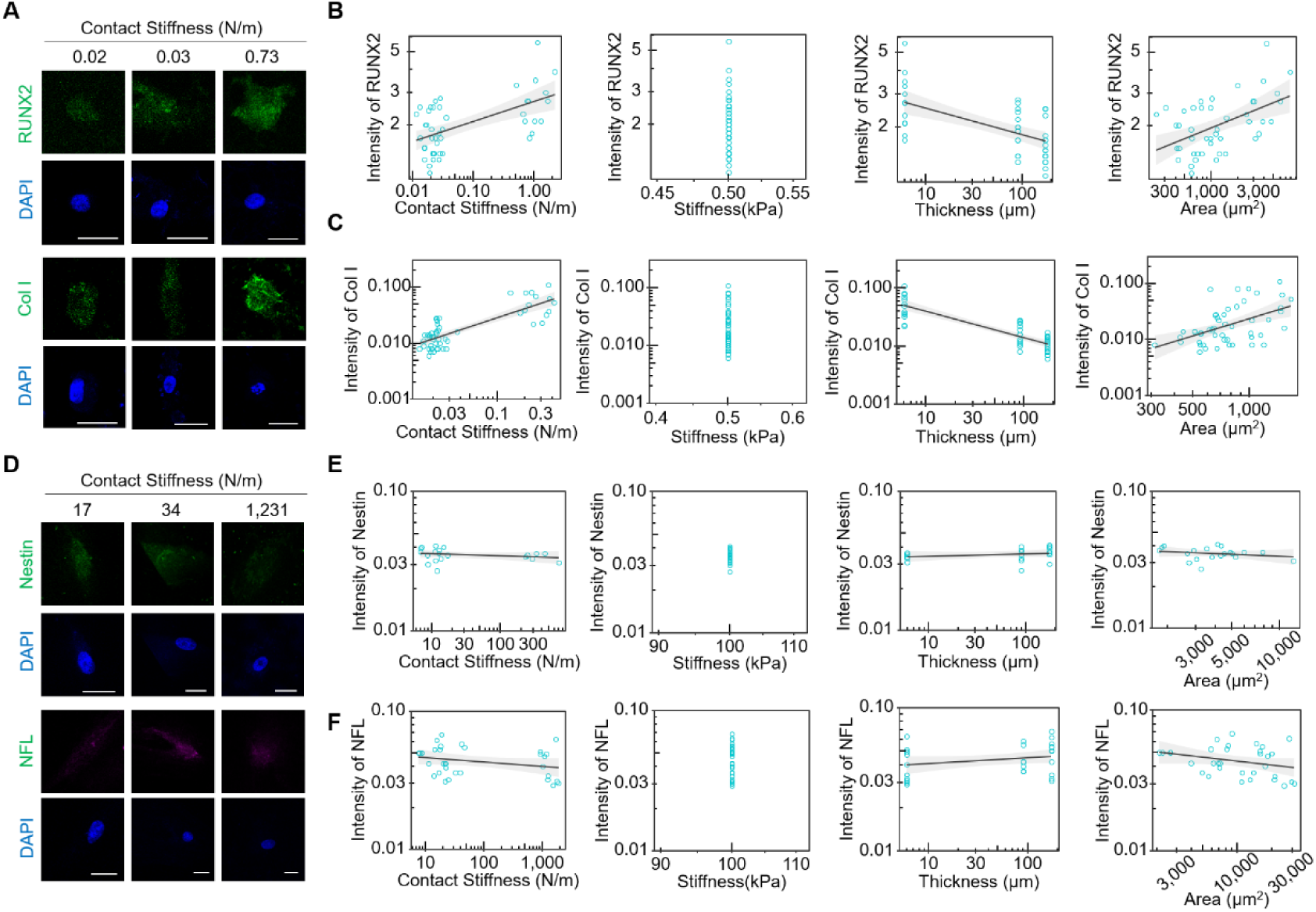
Relationship between stem cell differentiation marker levels and different variables of cell-ECM system. (**A**) Representative immunofluorescence images of Runx2, Collagen I (Col I) and nucleus in hMSCs grown on a 0.5 kPa gradient thickness gel for 4 days at different contact stiffness stained by Runx2 or Collagen I antibody (green) and DAPI (blue), respectively. Scale bar: 30 μm. (**B**) and (**C**) Relationship between (B) Runx2 (C) Collagen I expression level and contact stiffness or other parameters (ECM stiffness, ECM thickness and cell adhesion area) in hMSCs grown on a 0.5 kPa gradient thickness gel for 4 days. (B) RUNX2 level scales across all the datasets as power functions of Contact Stiffness, with exponent 0.116 ± 0.024 (*r^2^* = 0.335, *P* < 0.001); Thickness, with exponent −0.144 ± 0.031 (*r^2^* = 0.331, *P* < 0.001); Area, with exponent 0.238 ± 0.053 (*r^2^* = 0.291, *P* < 0.001); Grey area are 95% confidence intervals for the fitted functions. (C) Col I level scales across all the datasets as power functions of Contact Stiffness, with exponent 0.497 ± 0.07 (*r^2^*= 0.59, *P* < 0.001); Thickness, with exponent −0.455 ± 0.061 (*r^2^* = 0.65, *P* < 0.001); Area, with exponent 1.07 ± 0.304 (*r^2^* = 0.215, *P* < 0.001); Grey area are 95% confidence intervals for the fitted functions. (**D**) Representative immunofluorescence images of Nestin, Neurofilament (NFL) and nucleus in hMSCs grown on a 100 kPa gradient thickness gel for 4 days at different contact stiffness stained by Nestin or NFL antibody (green) and DAPI (blue), respectively. Scale bar: 30 μm. (E) and (F) Relationship between (**E**) Nestin (**F**) NFL level and contact stiffness or other parameters (ECM stiffness, ECM thickness and cell adhesion area) in hMSCs grown on a 100 kPa gradient thickness gel for 4 days. (E) Nestin level scales across all the datasets as power functions of Contact Stiffness, with exponent −0.015 ± 0.014 (*r^2^* = 0.057, *ns*, not significant); Thickness, with exponent −0.029 ± 0.021 (*r^2^* = 0.064, *ns*, not significant); Area, with exponent −0.052 ± 0.049 (*r^2^* = 0.059, *ns*, not significant); Grey area are 95% confidence intervals for the fitted functions. (F) NFL level scales across all the datasets as power functions of Contact Stiffness, with exponent −0.029 ± 0.021 (*r^2^* = 0.046, *ns*, not significant); Thickness, with exponent 0.042 ± 0.029 (*r^2^* = 0.063, *ns*, not significant); Area, with exponent −0.074 ± 0.053 (*r^2^*= 0.059, *ns*, not significant).

**Fig. S4.**
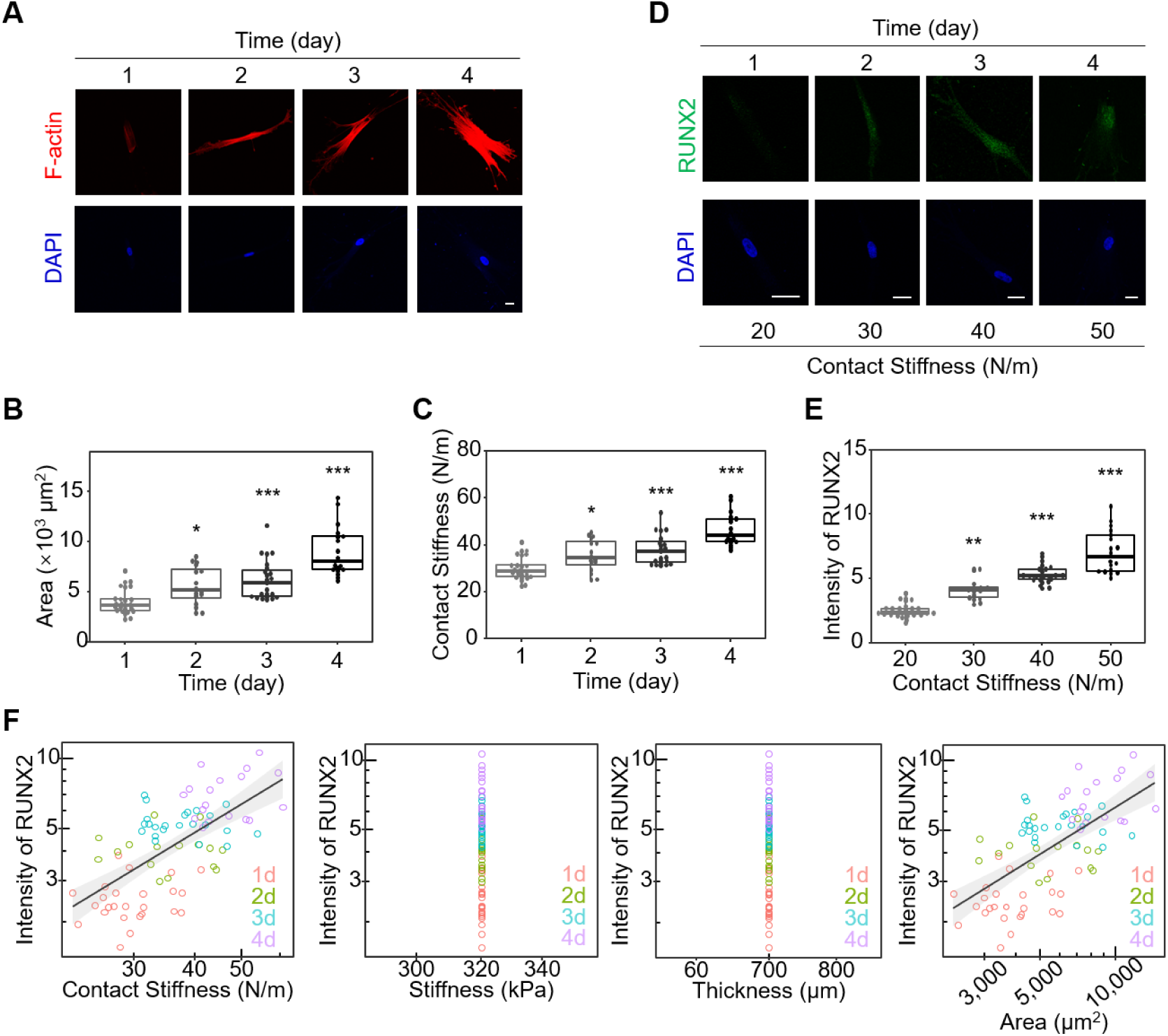
Contact stiffness scales with Runx2 level during stem cell differentiation induced by osteogenic differentiation medium. (**A**) Representative immunofluorescence images of F-actin and nucleus in hMSCs grown on a 320 kPa PDMS gel with constant thickness (larger than 200 μm) gel in the presence of osteogenic differentiation medium stained by phalloidin (red) and DAPI (blue), respectively. Scale bar: 30 μm. (**B**) The statistical analysis of cell adhesion area in hMSCs grown on a 320 kPa PDMS gel in the presence of osteogenic differentiation medium over time as shown in (A). All error bars are S.E.M. (1d: n = 25 cells; 2d: n = 16 cells; 3d: n = 23 cells; 4d: n = 18 cells. * *P* < 0.05; ** *P* < 0.01; *** *P* < 0.001; *ns*, not significant). (**C**) The statistical analysis of contact stiffness in hMSCs grown on a 320 kPa PDMS gel in the presence of osteogenic differentiation medium over time as shown in (A). All error bars are S.E.M. (1d: n = 25 cells; 2d: n = 16 cells; 3d: n = 23 cells; 4d: n = 18 cells. * *P* < 0.05; ** *P* < 0.01; *** *P* < 0.001; *ns*, not significant). (**D**) Representative immunofluorescence images of RUNX2 and nucleus in hMSCs grown on a 320 kPa PDMS gel with constant thickness (larger than 200 μm) gel in the presence of osteogenic differentiation medium stained by RUNX2 antibody (green) and DAPI (blue), respectively. Scale bar: 30 μm. (**E**) The statistical analysis of RUNX2 intensity in hMSCs grown on a 320 kPa PDMS gel in the presence of osteogenic differentiation medium over time as shown in (A). All error bars are S.E.M. (n = 25, 16, 23, and 18 cells for CS = 20, 30, 40, and 50 N/m, respectively. * *P* < 0.05; ** *P* < 0.01; *** *P* < 0.001; *ns*, not significant). (**F**) Relationship between RUNX2 expression level and contact stiffness or other parameters (ECM stiffness, ECM thickness and cell adhesion area) in hMSCs grown on a 100 kPa PDMS gel in the presence of osteogenic differentiation medium. RUNX2 level scales across all the datasets as power functions of Contact Stiffness, with exponent 1.16 ± 0.159 (*r^2^* = 0.408, *P* < 0.001); Area, with exponent 0.627 ± 0.086 (*r^2^* = 0.407, *P* < 0.001); Grey area are 95% confidence intervals for the fitted functions.

**Fig. S5.**
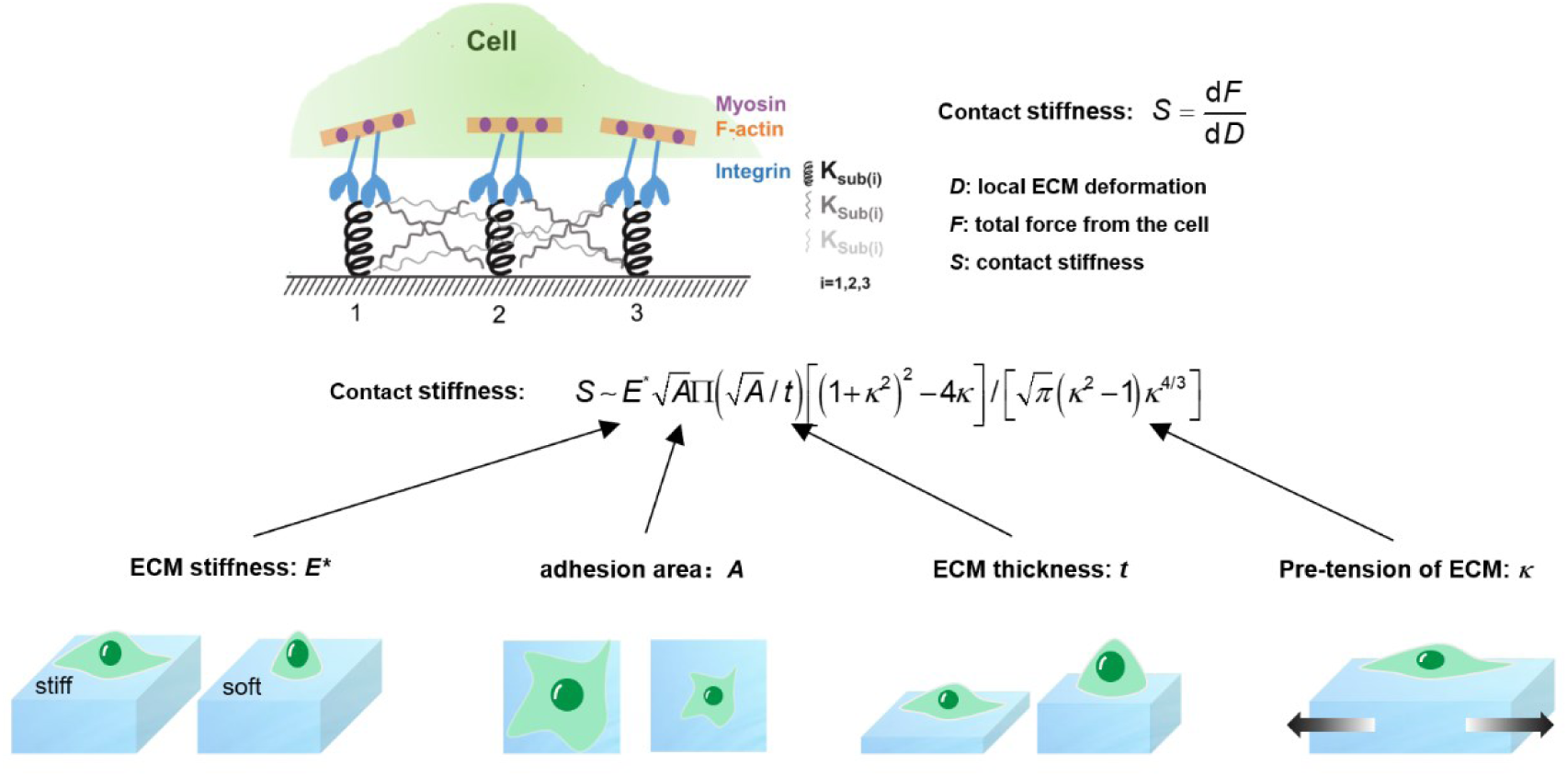
Schematic diagram of a CS-based frame of reference for interpreting the effects of various ECM mechanics on cell behaviors. CS defines the relationship between local ECM deformation and the force imposed by the cell, integrating mechanical variables such as ECM stiffness (as measured by the elastic and viscoelastic properties), ECM thickness, pre-tension of the ECM, and cell adhesion area into one variable.

**Fig. S6.**
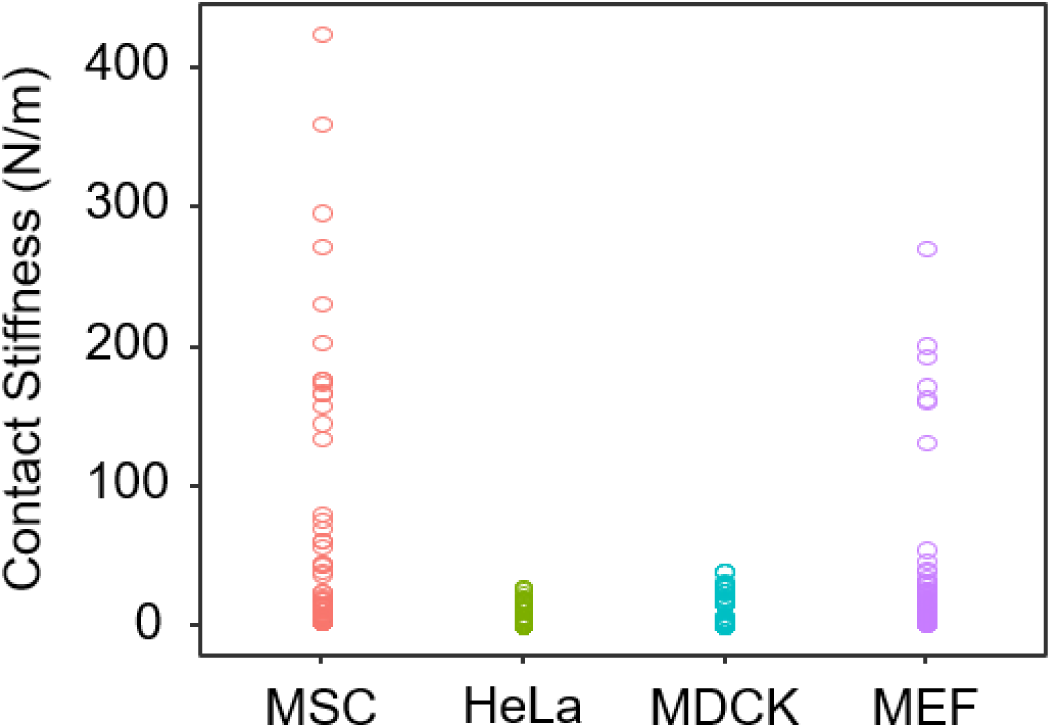
The variation range of the CS in different types of cells. The statistical analysis of CS in different types of cells (hMSCs, HeLa cells, MDCK cells and MEFs) grown on 40 kPa gradient thickness gels.

**Fig. S7.**
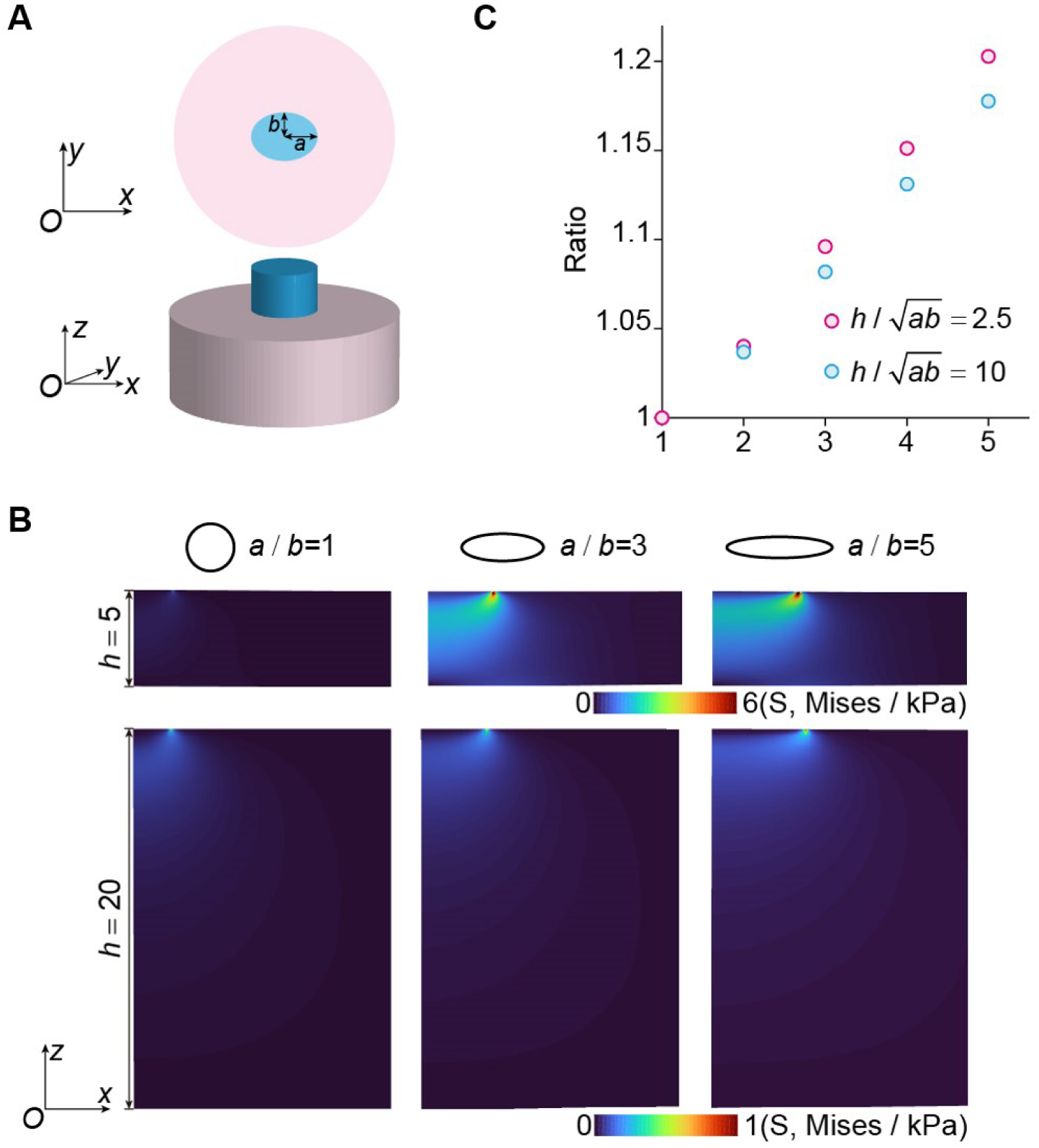
Finite element analysis of the effect contact geometry on the contact stiffness (the contact areas are the same for all the cases). (**A**) Finite element model of the contact with elliptical contact region. (**B**) Contour of stresses in the contact region for different contact geometries and substrate thickness. The results show that the substrate effect will be significant when the substrate is thin and the ratio of a/b is large. (**C**) The ratios of the contact stiffness for different contact geometries with respect to that corresponding to circular contact region.

